# Ethological variability shapes the structure and content of sensory population codes

**DOI:** 10.64898/2026.10.13.758815

**Authors:** Augustine (Xiaoran) Yuan, Hongyu Li, Steffen Katzner, Laura Busse, Wiktor Młynarski

## Abstract

Established theories of sensory coding posit that sensory systems are adapted to the statistics of natural stimuli that originate in the external environment. In natural, ethological conditions, however, sensory populations are simultaneously influenced by multiple modulatory processes. During behavior, the statistics of population responses already at the sensory periphery reflect not only external signals, but also internal and behavioral states such as locomotion, attention, and arousal. Theories of sensory coding in real-world conditions must therefore explain adaptation not only to natural scenes, but also to the structured variability induced by these internal processes and behavior. Here, we investigated population activity in the dorsal lateral geniculate nucleus (dLGN), a critical early stage of the visual pathway. Using a simple probabilistic model and information-theoretic approaches, we characterized this variability through the statistics of the population messages conveyed downstream. Our results show that population variability is multidimensional: even when one behavioral factor is constant, responses can vary substantially as other modulatory factors fluctuate. Under natural conditions, sensory population codes can therefore change their statistical structure and content, i.e., the stimulus features they encode along multiple dimensions simultaneously. These findings highlight the need for sensory coding theories that account not only for adaptation to natural stimuli, but also to the statistics of internal and behavioral states.

## INTRODUCTION

Our theoretical understanding of sensory systems has long been grounded in the stimulus-centric framework, where neurons at progressive stages of the sensory hierarchy extract and encode increasingly more complex features of the sensory input [1, 2]. In this traditional view, the statistics of sensory messages conveyed from the periphery to the cortex, as well as the computations performed at different stages of sensory processing, are assumed to depend predominantly on the physical properties of the external environment. Contemporary experiments in awake behaving animals reveal a crucial, previously overlooked aspect of sensory coding: the activity of neural populations is only partially influenced by external signals, even at the early processing stages and when the sensory input remains constant. Instead, sensory responses are impacted and modulated by many additional factors. Some of them correspond to overtly manifested behavior, e.g., running speed [3] or pupil area fluctuations [4, 5]. Other factors that contribute to neural variability are more internal, including attention [6] or task engagement [7]. Each of these factors could, in principle, correspond to many modulatory variables that simultaneously influence the sensory code over multiple timescales [5]. These internal states can also modulate and interact with sensory representations through spontaneous activity [8, 9]. Recent evidence demonstrates that such modulation applies not only to neurons in the visual cortex [10], but extends through the thalamus all the way to the sensory periphery [11–14], implying that even direct “measurements” of the sensory environment are significantly affected by a multitude of internal and behavioral states. Understanding sensory coding during natural behavior therefore requires approaches that quantify and identify structure in such variability.

These experimental developments present a challenge to existing theories of sensory coding. The efficient coding hypothesis — a conceptual cornerstone of sensory neuroscience — postulates that the brain adapts to the statistical structure of natural stimuli encountered by animals in their ecological niche [1, 15]. This adaptation may extend from the sensory periphery [16–19] to higher stages of visual processing [20–23]. Efficient coding theory has been extended to postulate different objectives of sensory coding, such as prediction of the future input [24, 25] or inference of latent variables [26]. Other extensions accentuate the dynamic nature of sensory inputs and the necessity for sensory adaptation as a way to maintain efficient codes in changing environments [27– 29]. Despite these developments, the dominant theory of sensory systems remains stimulus-centric: the central assumption is that “messages” conveyed by neural populations up the sensory hierarchy are determined by the external stimulus. In reality, however, the statistics of natural sensory messages are heavily shaped by active behavior, changing objectives, and simultaneously fluctuating internal states. To understand sensory processing during natural behavior, we must embrace the complexity of such dynamic and nonstationary sensory codes.

In this work, we take steps toward extending the efficient coding framework to account for the statistics of natural sensory messages rather than solely natural stimuli. We study neural populations in the dorsal lateral geniculate nucleus (dLGN) of the mouse — a key stage of the early visual processing. Neurons in dLGN receive direct input from the retina — the retinal ganglion cells (RGCs). At the same time, they receive descending inputs from the visual cortex and other brain areas [30]. Thalamic areas such as dLGN are also subject to rich behavioral modulation by running speed [11] and internal states, manifested by pupil area dynamics [5]. Neural populations in the dLGN are therefore an ideal model system to study the impact of multiple modulatory factors on the statistics of sensory messages sent downstream, along the hierarchy of the visual pathway.

To understand variation of the sensory codes, we developed a simple probabilistic model of population activity patterns at the level of individual spikes. Despite its simplicity, the model was able to capture the nonstationary nature of neural population dynamics. We then used the model fits to characterize each experimental trial in terms of the statistics of population-level messages sent downstream. We show that the population codes vary continuously along multiple dimensions. Crucially, we demonstrate that variation along these dimensions changes the statistical structure of the population code, as well as its meaning, i.e., the stimulus features it represents.

Our approach provides a conceptual and quantitative framework to characterize statistics of sensory messages conveyed by neural populations during natural behavior. Our results also highlight the need for theories of sensory systems grounded not only in the statistics of natural scenes, but also in the structure of natural sensory messages impacted by varying states of the behaving organism.

## RESULTS

### Population codes in the thalamus are highly variable during behavior

Populations of sensory neurons process and encode not only signals from the environment, but also from signals inside the nervous system (Fig. 1a). These internal signals can be roughly assigned to two broad classes. The first class consists of signals correlated with the overt behavior of the animal, such as running speed or changes in the pupil area [3]. The second class consists of signals that are purely internal and may be difficult or impossible to read out from the outside [31].

**Figure 1:**
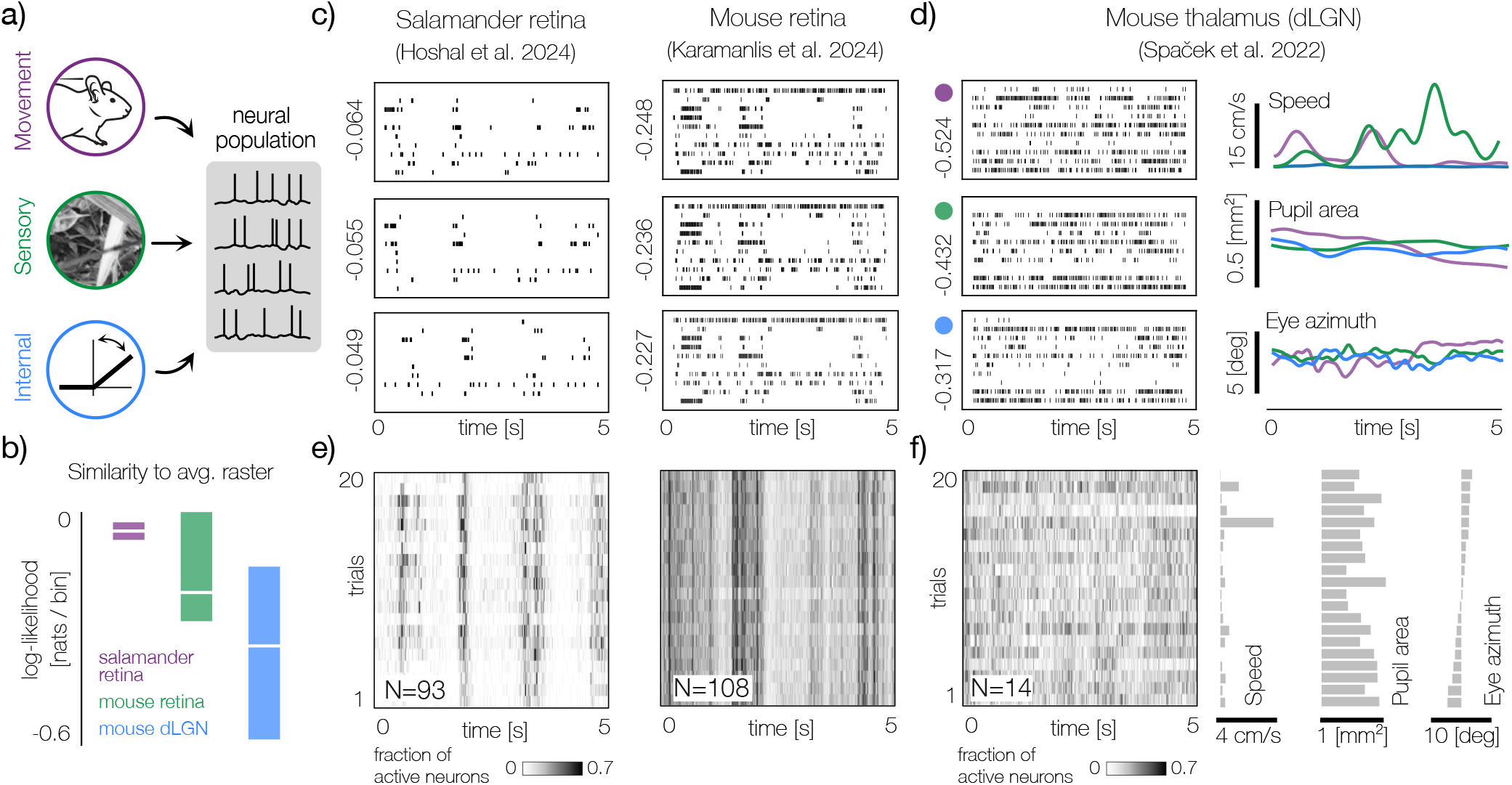
Variability of population codes in dLGN during behavior. **a)** Neural populations in the thalamus are driven by stimuli (green), movement of the animal (purple) and internal modulatory processes (blue). **b)** Distributions of log-likelihood values of individual trials under the average raster model (see Methods) for neural populations in the salamander retina (taken from [32]; purple), mouse retina (taken from [33]; green) and mouse dLGN (blue). White lines denote medians, rectangles span the range between 1st and 99th percentiles. **c)** Left: responses of a population of 10 salamander retinal ganglion cells (RGCs) to three presentations of a natural video (data from [32]). Numbers correspond to values of log-likelihood displayed in b). Right: the same for a population of mouse RGCs (data from [33]). **d)** Left: responses of a population of 10 neurons in the mouse dLGN to three presentations of a natural video recorded during behavior in a head-fixed setup (data from [30]). Right: Trajectories of behavioral variables. Lines correspond to trials marked with the same color in the left column. **e)** Population synchrony, i.e. the fraction of active neurons in the population during presentation of a stimulus over 20 trials. Left: population of 93 RGCs in the salamander retina. Right: population of 108 RGCs in the mouse retina. **f)** Left: population synchrony in a population of 14 neurons in the mouse dGLN over 20 trials. Right: average values of behavioral parameters (running speed, pupil area and azimuthal eye position) on each trial. Trials are sorted by the eye azimuth.

To illustrate how these factors contribute to the variability of population responses already at the early stages of the visual pathway, we analyze the activity of a small neural population recorded in the dLGN of a behaving mouse (see Methods; data published previously in [30]). We contrast this recording with populations of retinal ganglion cells recorded ex-vivo in the salamander (data from [32]) and mouse (data from [33]). In all cases, neurons were exposed to short clips of natural videos that may approximate ecologically valid stimuli.

Population responses in the retina are largely reproducible across trials (Fig. 1c). Despite non-trivial and highly structured stochasticity, which is famously non-trivial and highly structured (see, e.g., [34–37]), retinal responses to the same stimulus are reminiscent of each other. At a population level, this effect can be summarized by displaying synchrony, i.e., the fraction of coactive neurons in any given time bin and trial (Fig. 1e). While some variability is visible, the patterns of co-activation are similar across repeated exposures to the identical stimulus.

Recordings in the ex-vivo retina can be interpreted as an approximation of direct inputs to the dLGN (although we note that retinal outputs are already modulated during behavior [12], [38]). In contrast to the reproducibility of RGC responses, population activity in the dLGN of a behaving mouse can be highly variable (Fig. 1d, left column). At first glimpse, example population rasters seem to be entirely different, even though the eye position (and therefore the sensory input) is practically identical on each trial (Fig. 1d, right column, bottom row). The extent of trial-to-trial variability is visible in the synchrony plot of the dLGN population (Fig. 1f, left panel).

To quantify this variability directly, we introduced a simple, coarse metric of similarity of a population raster on a single trial to the trial average (Fig.1b; see Methods). The difference between the ex-vivo retina and the dLGN during behavior is clearly very strong. Not only is the mean similarity of individual trials to the average population response much smaller across the analyzed dLGN populations than in the retinal datasets, but the variance of the similarity measure is also substantially higher in the dLGN. The analysis therefore demonstrates that during behavior, sensory codes become non-stationary and highly variable already one synapse away from the retina. Regardless of the function and origin of these changes, downstream circuits must be able to adapt to their structure in order to efficiently process inputs from dLGN and flexibly perform different computations.

### A simple probabilistic model captures the population variability

To characterize population activity at the single-spike level, we represent the population state in each 20 ms bin by a binary vector 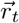, where entry *n* equals 1 if the corresponding neuron spikes in that bin and 0 otherwise. Because the number of possible spike patterns grows exponentially with population size, quantifying dLGN coding during behavior requires a statistical model that captures the essential structure of population activity [34, 35, 37, 39].

Population models have yielded major advances in sensory coding, but most assume stationarity (i.e., time-invariant statistics). dLGN codes violate this assumption: population statistics evolve over the experiment, and individual trials may be drawn from different underlying distributions. As a result, we must characterize an exponentially large state space while estimating parameters that vary over time, often from limited samples.

To address this challenge, we use a latent-variable approach [40]. Latent-variable models describe population activity using a small set of shared factors that capture joint firing patterns. Because these factors can evolve over time, they naturally capture non-stationary population dynamics while remaining constrained by correlations and frequently co-occurring activity patterns. Related approaches have also modeled population activity at the level of individual spikes, for example, using restricted Boltzmann machines [36, 41, 42].

Building on these ideas, we assume that each neuron’s spiking probability at each time point is controlled by a set of latent variables whose values evolve over time. Each latent variable is associated with a weight vector that determines whether it excites or inhibits each neuron, thereby increasing or decreasing its firing probability (Fig. 2a,b). The model thus decomposes the rasters of population activity into latent variable activations and their associated weights; we therefore call it probabilistic raster factorization (PRF). We note, however, that it is closely related to logistic matrix factorization [43], with the only distinction being that the bias terms are set to match the baseline spiking probability of each neuron (see Methods). It also bears resemblance to the restricted Boltzmann machine (RBM) [42]. The difference, however, is that the latent variables in the RBM are binary. Due to its simplicity, the PRF model can be readily fitted to the data by maximizing the log-posterior using gradient-based approaches (see Methods). During fitting, we apply a prior that promotes minimal deviation of each neuron from its baseline activity (see Methods).

**Figure 2:**
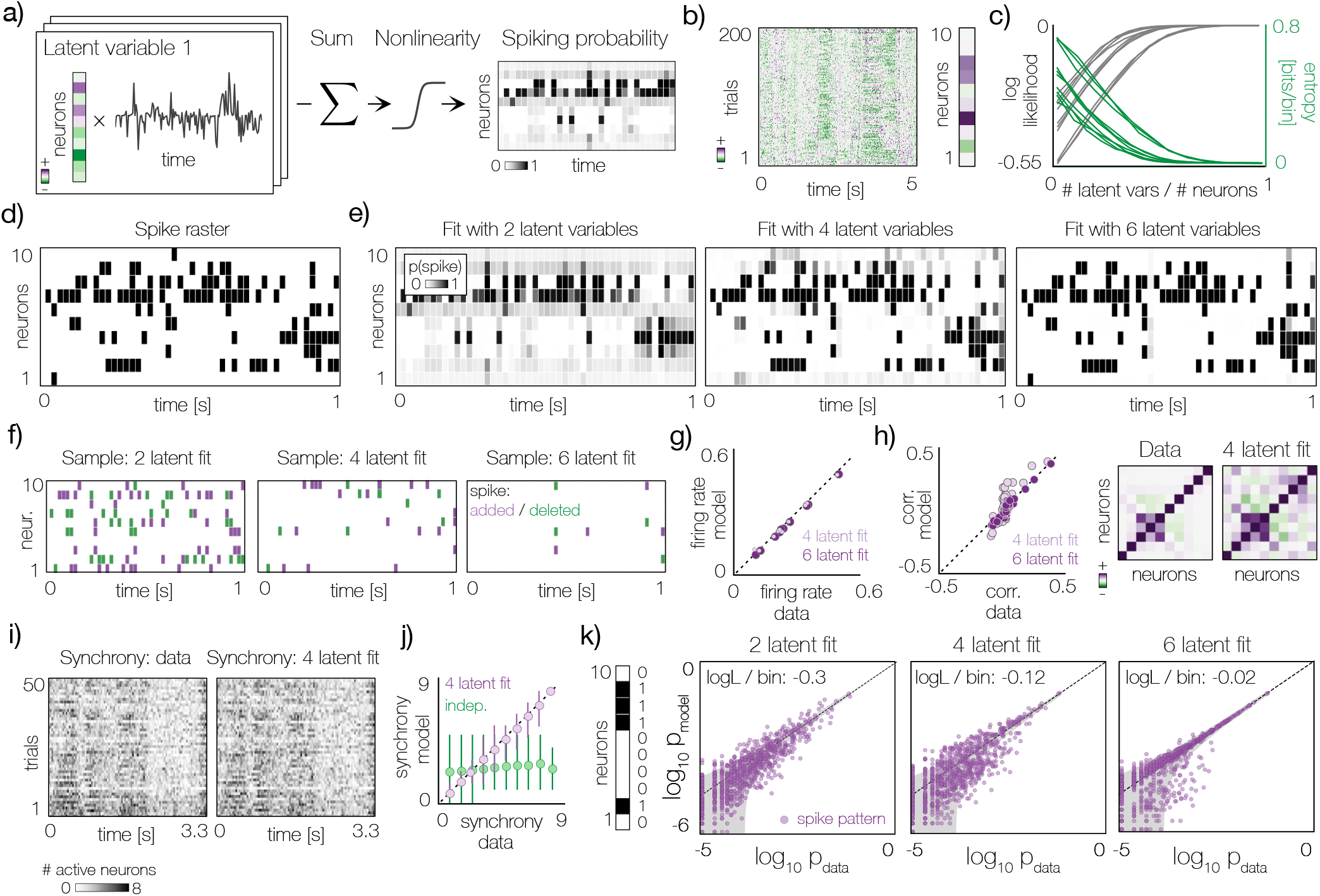
Probabilistic Raster Factorization model captures nonstationary statistics of population activity. **a)** Schematic of the PRF model. Spiking probability patterns are decomposed into activity of a set of latent variables. **b)** Example latent variable across trials (left) controls firing probabilities of neurons within the population (right). **c)** Gray lines: data log-likelihood vs number of latent variables relative to the number of neurons. Each line corresponds to an individual dataset. Green lines: entropy per bin vs the relative number of latent variables. **d)** Example excerpt of a population raster. **e)** Spiking probabilities corresponding to the excerpt in d. From left to right - models with increasing number of latent variables. **f)** Population rasters sampled from the model fits visualized in e. Purple: spikes that do not exist in the original raster (panel d) but do exist in the sample. Green: spikes that exist in the original raster (panel d) but do not exist in the sample. **g)** Average firing rates vs empirical firing rates for two model fits with 4 (small circles, light purple) and 6 (large circles, dark purple) latent variables. **h)** Left: pairwise correlations in the data and obtained from model samples. Middle: empirical correlation matrix. Right: correlation matrix obtained from samples from a model fit with 4 latent variables. **i)** Left: synchrony values across 50 trials in the data. Right: synchrony in the same trials averaged across samples from a model fit with 4 latent variables. **j)** Empirical synchrony values vs synchrony values obtained from model samples. Circles correspond to averages, lines to the interval between 5th and 95th percentiles. Purple: PRF model fit with 4 latent variables. Green: Samples from independent model. **k)** Empirical probability of population codewords (purple circles; illustrated in the left column) vs probability obtained from sampling PRF model fits with increasing numbers of latent variables. Gray funnels correspond to 99th confidence intervals.

The key parameter of the PRF model is the number of latent variables. If there are as many of them as neurons, each individual spike and silence is explained perfectly and assigned a likelihood of 1. When no latent variables are fitted, the probability of each neuron spiking is equal to the baseline firing rate across all time bins, which determines the minimal likelihood value. Any number of latent variables between these two extremes interpolates between “perfect explanation” of the data and the baseline firing rate, where firing probabilities are constant (Fig. 2c, gray lines).

Complementary to the likelihood, we can consider the uncertainty about the firing of each neuron, which is measured by the binary entropy of the spiking probability in each bin (Fig. 2c, green lines; see Methods). The total amount of uncertainty to be explained by the model is determined by the baseline firing probabilities. The objective of the PRF model is to “explain away” this uncertainty. If the number of latent variables is equal to the number of neurons, the uncertainty about each spike or silence vanishes. This approach is conceptually related to principal component analysis (PCA) - a standard dimensionality reduction method. PCA identifies components (i.e., latent variables) that explain the total variance of the data. This approach therefore enables us to retain the set of components that capture significant trends in the data and do not represent noise and random fluctuations.

In population spiking data, such a decomposition corresponds to identifying patterns of activity that systematically co-occur across multiple neurons and may therefore reflect shared inputs and co-modulation. Importantly, individual spikes are stochastic; therefore, two individual population responses may not be exactly identical, even when the underlying inputs are identical. We illustrate this intuition by fitting the PRF model to a population of 10 neurons and selecting a single 1-second excerpt of activity (Fig. 2d). While the PRF model with two latent variables already identifies significant patterns of co-firing and assigns them high probability, a lot of uncertainty about activity remains (Fig. 2e, left panel). Increasing the number of latent variables to six dramatically improves the quality of the fit and reduces the uncertainty about the individual spikes and silences.

In our analyses, we need to set an accuracy threshold at which we will consider that any residual uncertainty is due to the inherent stochasticity of spiking, rather than to significant inputs and modulatory influences. That threshold, in turn, determines the number of latent variables that capture the non-stationary distribution of population activity. Here, we set the accuracy threshold of log-likelihood such that the probability of each spike or silence given the model fit is not smaller than −0.1 (see Methods). Given this criterion, the dimensionality of the relatively small neural populations analyzed here can be reduced by 50–70 % with respect to the number of neurons in the population (Fig. 2c).

The fitted PRF model specifies a non-stationary distribution over the population activity. This means that we can generate samples of artificial population activity consistent with the fit (Fig. 2f). Following the established methodology, we then test how closely do these samples reproduce the statistics of the data that were not directly fitted by the model [34, 44]. We note that, as in the RBM model, neuron spiking probabilities are conditionally independent in each time bin *t* given the state of the latent variables and associated weights. The existence of temporal correlations between latent variables (both within and across trials) introduces dependencies between neurons. Samples from the PRF model fit can capture correlations and their non-stationary structure present in the data.

We find that sample statistics closely match marginal statistics such as firing rates (determined by spiking probabilities) and neural correlations (Fig. 2g–h). To further verify how well the model fits the temporal structure of population activity, we examine synchrony across individual trials (Fig. 2i). Model samples closely match this feature of the data and substantially outperform an independent model (Fig. 2j). To further confirm the goodness of fit, we compared empirical probabilities of spike patterns with those obtained from model samples and obtained a reasonable match between the two (Fig. 2k). Altogether, these analyses indicate that the PRF model can accurately characterize the structure of population activity at the level of individual spikes.

### Latent variable statistics reveal nonstationary properties of the neural code

Visualizing latent variables across trials reveals a non-stationary structure in population activity (Fig. 3a). This nonstationarity is captured by the time-averaged latent variables on each trial (Fig. 3b): some averages exhibit slow drift across trials, whereas others show more abrupt fluctuations. In a stationary code, these averages would be distributed around a stable mean, which is clearly not the case here.

**Figure 3:**
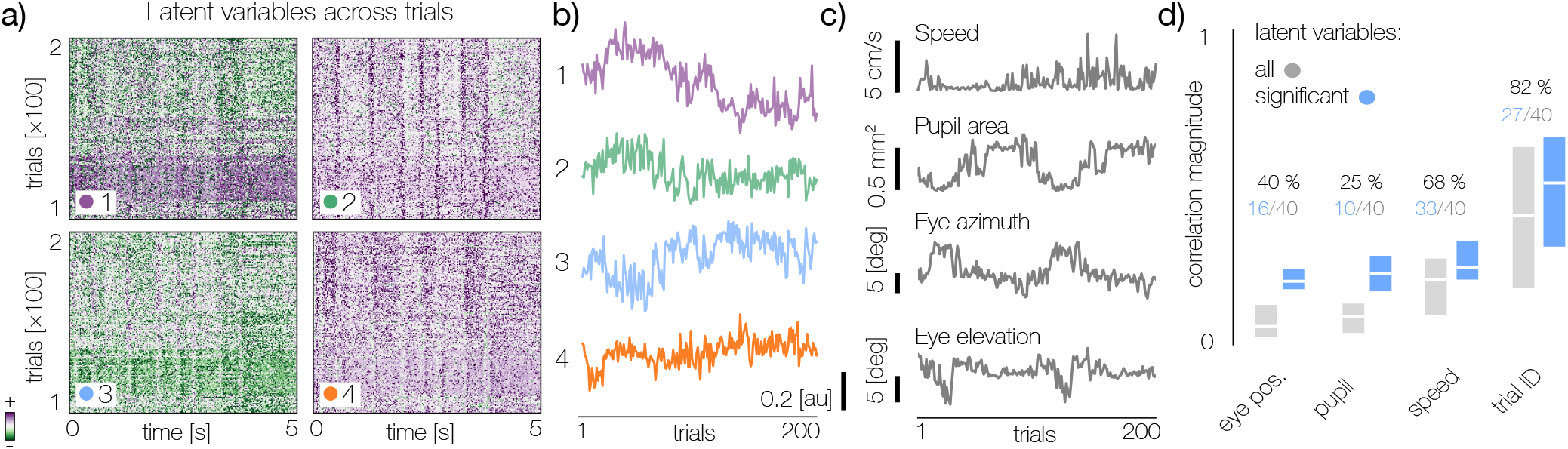
Statistics of latent variables reveal nonstationary population dynamics. **a)** Trajectories of four latent variables across trials. The latent variables were fitted to a population of 20 neurons. **b)** Time-averages of latent variables reveal changing statistics of population activity. Line colors correspond to color markers in a). **c)** Time averages of behavioral variables across trials. **d)** Distributions of correlation magnitudes (absolute values) between trial-averages of latent and behavioral variables, pooled across 9 analyzed populations. Bars denote the 25th-75th percentile intervals and horizontal white lines denote median. Gray bars denote all correlations, blue bars denote only significantly correlated latent variables (determined by a permutation test - see Methods).

We compared trial-to-trial changes in the time-averaged latent variables with behavioral parameters, including running speed, pupil area, and eye position (Fig. 3c). Although changes in these parameters are different from changes in the latent variables, they evolve qualitatively similarly over the experiment, with both gradual drift and sudden shifts. As with latent-variable models in general [45], the PRF model does not guarantee a clean separation of the biological processes underlying non-stationary population dynamics. Nevertheless, individual latent variables, while not directly interpretable, can be meaningfully related to behavior via correlation analyses (Fig. 3d).

Across datasets, changes in the statistical structure of the code are most strongly correlated with time in the experiment, as indexed by trial number. This suggests that slow fluctuations in the organism’s internal state may exert the largest influence on sensory representations. Running speed shows the next-strongest correlation, followed by pupil area. By contrast, mean eye position changes in azimuth and elevation are only weakly correlated with the latent variables. Thus, although sensory inputs may vary with trial-to-trial eye movements, this variation appears substantially smaller than the variability driven by internal modulators.

### Multidimensional variability of population codes

The presence of multiple modulators that simultaneously shape dLGN population activity has important consequences for downstream areas. Under ethological conditions, neurons in the primary visual cortex may receive spike patterns from dLGN populations that follow very different distributions, even when the sensory input itself is unchanged (Fig. 4a). A natural way to quantify the difference between these distributions is the Jensen-Shannon divergence (*D*_JS_; see Methods), an information-theoretic dissimilarity measure that is equal to 0 bits when two distributions are identical and 1 bit when they are entirely non-overlapping (Fig. 4b). We leverage the PRF model to estimate the distributions of spike patterns conveyed by the dLGN on each trial and compute the corresponding *D*_JS_ between trials (Fig. 4c).

**Figure 4:**
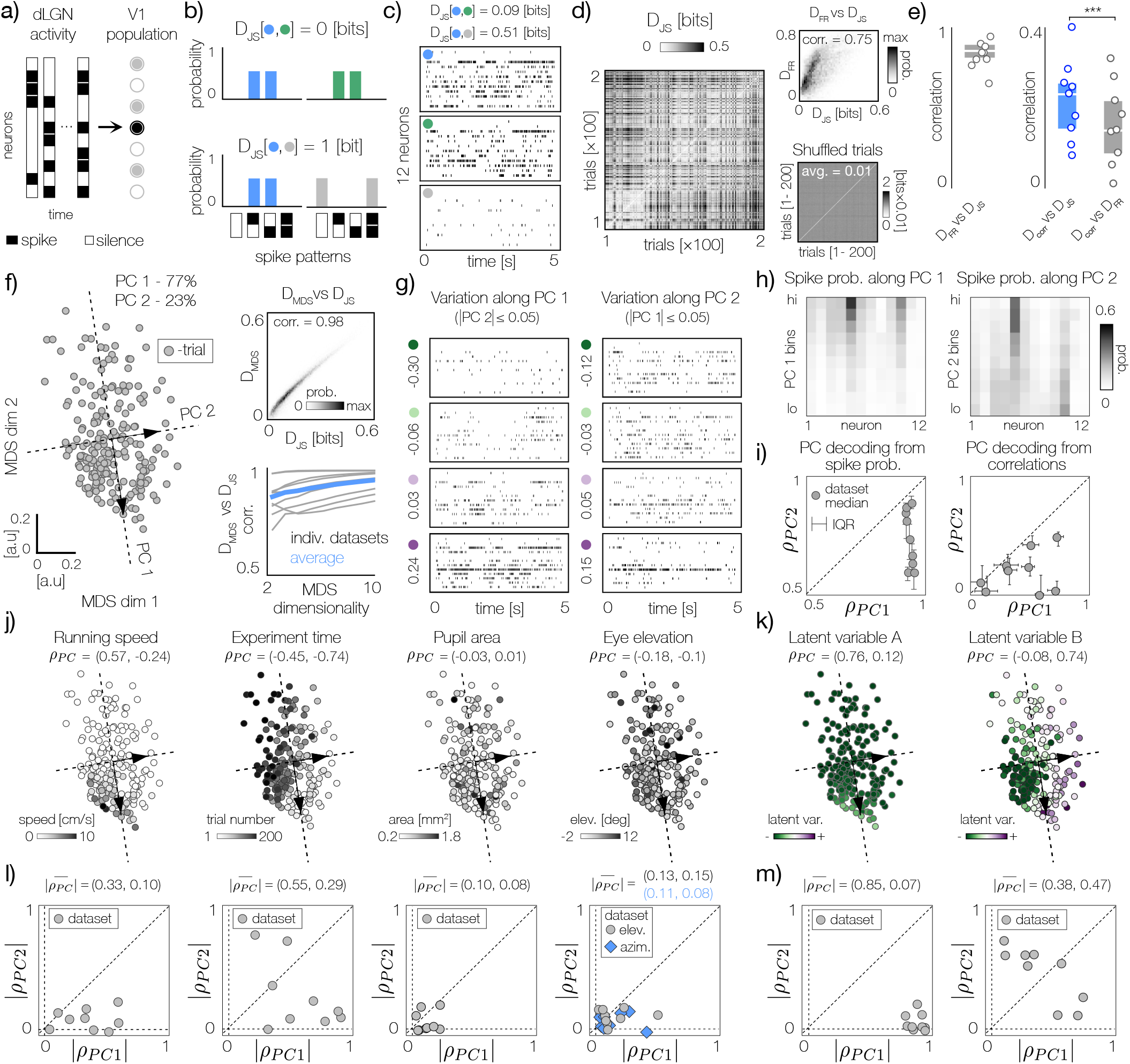
Multidimensional variability of dLGN population codes. **a)** Downstream areas (e.g. V1) process population-level spike patterns generated by dLGN. **b)** Jensen-Shannon divergence (*D*_*JS*_) measures the difference between spike pattern distributions. **c)** Top: example reference trial of a population of 12 neurons. Middle: trial most similar to the reference as quantified by *D*_*JS*_. Bottom: trial most different from the reference. **d)** Left: a matrix of *D*_*JS*_ distance between all trials in a population of 12 neurons. Right bottom: same as left but for randomized trials with inter-trial differences removed. Right top: joint distribution and correlation of *D*_*JS*_ and Euclidean distance between firing rates *D*_*F R*_. **e)** Left: correlations of *D*_*JS*_ and *D*_*F R*_ between trials. Each dot corresponds to a dataset. Gray bar and white lines correspond to the 25th-75th percentile range and the median respectively. Right: blue - correlations of *D*_*JS*_ between trials and Euclidean distance between correlation matrices *D*_*corr*_. Gray - correlations of *D*_*corr*_ and *D*_*F R*_. All markers as in the left panel. Stars denote statistically significant difference (paired sample t-test, *t* = 7.051, p-value = 1.07 × 10^−4^). **f)** Left: two-dimensional MDS projection of 200 trials based on their distance. Gray circles denote individual trials. Dashed lines and vectors denote two principal components. Right bottom: correlation between *D*_*JS*_ and Euclidean distance between MDS projections *D*_*MDS*_ as a function of projection dimensionality. Gray lines denote individual datasets and the blue line the average. Right top: joint distribution and correlation of *D*_*JS*_ and Euclidean distance between firing rates *D*_*MDS*_. **g)** Left: raster plots sorted along the first principal component (PC 1) with the second component close to 0. Values of the projection on PC 1 are displayed on the side of each raster. Right: the same as left but for the second principal component (PC 2). **h)** Left: change of spiking probability along PC 1. The range of coordinates along PC 1 was divided into 10 bins sorted from the lowest to highest values. Right: the same as left but for PC 2. **i)** Dataset summary - decoding of PC 1 and PC 2 coordinates from neural activity across all datasets. Left: decoding from trial-average spiking probability (firing rate). Correlation of the decoded coordinate along PC 1 with the true value (horizontal axis, *ρ*_*P C*1_) plotted against the correlation of the decoded PC 2 coordinate with the true value (vertical axis, *ρ*_*P C*2_). Each symbol corresponds to an individual dataset. Circles denote median correlation among random splits, bars the interval between 25th - 75th percentile. Right: same as left but for decoding of PC coordinates from correlations on each trial. **j)** MDS projections of individual trials (as in the panel to f)). Grayscale colors correspond to average value per trial of different variables averaged across trial duration. From left to right: running speed, experiment time, pupil area and eye elevation. *ρ*_*P C*_ denotes a pair of correlation values between the average variable and the first and second principal component respectively. **k)** Same as j) but colors correspond to time averages of the two latent variables A and B that are most strongly correlated with PC 1 and PC 2 coordinates, respectively. **l)** Dataset summary - correlation magnitudes with PC 1 and PC 2 (|*ρ*_*P C*1_|, |*ρ*_*P C*2_|). Each panel corresponds to the variable plotted above in k). Dots corresponds to datasets. Blue diamonds in the rightmost panel denote eye azimuth plotted in addition to eye elevation. **m)** Dataset summary. Same as l) but for latent variables most strongly correlated with PC 1 and PC 2 coordinates respectively.

The *D*_JS_ dissimilarity matrix reveals significant variability in spike pattern distributions across trials (Fig. 4d, left). This variability is not artifactual: when computed on shuffled data with stationary trial statistics, *D*_JS_ values effectively disappear (Fig. 4d, right column, bottom panel). To understand which features of neural activity contribute to distributional differences, we computed the Euclidean distance between vectors of spiking probabilities for each trial (*D*_FR_). The spiking-probability distance is strongly correlated with *D*_JS_ in the example population displayed in Fig. 4d, right top, as well as across all analyzed datasets (Fig. 4e, left), suggesting that gain changes in individualneuron firing rates contribute substantially to the non-stationary structure of the population code. At the same time, distances between correlation matrices on each trial (*D*_corr_) correlate more strongly with *D*_JS_ than with *D*_FR_, implying that information-theoretic dissimilarity captures aspects of spike pattern distributions that go beyond simple changes in spiking probability.

The dissimilarity structure in Fig. 4d suggests continuous variability in spike pattern distributions. To characterize this structure, we adopt a manifold learning approach based on multidimensional scaling (MDS). MDS is an embedding algorithm that projects high-dimensional spike pattern distributions onto a lower-dimensional representation while preserving the distances between distributions (quantified here by *D*_JS_ dissimilarity). Figure 4f displays a two-dimensional embedding generated from the dissimilarity matrix in Fig. 4d. In this representation, each point corresponds to a distribution of spike patterns generated by the population on a given trial, and the distance between points approximates the *D*_JS_ dissimilarity. The Euclidean distance on the MDS plane (denoted here by *D*_MDS_) strongly correlates with *D*_JS_ in the example population shown in Fig. 4f (right column, top panel). This agreement suggests that the code supported by this population varies primarily along two dimensions. In other analyzed populations, however, more dimensions may be required to accurately approximate *D*_JS_ between spike pattern distributions (Fig. 4f, right column, bottom panel). Although this implies that the population code can vary across multiple dimensions, for simplicity, we focus on a two-dimensional analysis of the studied populations.

The cloud of points generated by the MDS embedding is arbitrarily rotated. To parameterize trials in a more natural coordinate system, we performed principal component analysis (PCA) to identify directions of maximal variance among the spike pattern distributions on the MDS plane (Fig. 4f, black arrows and dashed lines). Raster plots of trials sorted along one principal component (PC), with the other held near zero, change systematically, with the nature of that change depending on which PC varies (Fig. 4g). In the example population in Fig. 4, movement along the first PC produces an overall increase in individual-neuron firing rates (Fig. 4h, left panel), whereas movement along the second PC produces a more nuanced shift in spiking probabilities, with some neurons increasing activity and others becoming quiet (Fig. 4h, right panel; e.g., neurons 5 and 10). To demonstrate that such changes occur systematically across all analyzed datasets, we decoded the PC coordinates from time-averaged spiking probability vectors on each trial (Fig. 4i, left panel). We found that the first PC can be reliably decoded across datasets, and decoding accuracy for the second PC is only slightly lower, as quantified by the correlation between decoded and ground-truth values (*ρ*_PC1_ and *ρ*_PC2_). Decoding PC coordinates from trial-wise correlation matrices is substantially less reliable, though it reaches relatively high values in some populations. This suggests that the population state may be most reliably identified based on simple features of population activity such as firing probabilities, rather than second order statistics such as correlations. Taken together, these decoding analyses demonstrate the generality of our approach across datasets and suggest that downstream circuits could reliably infer the state of the population code using simple activity statistics, even when the code is modulated by multiple concurrent factors.

The manifold analysis provides an effective visual representation of how the population code is simultaneously modulated by multiple factors by coloring the points representing each trial according to experimental and behavioral variables (Fig. 4j). To quantify these effects, directions of variation in spike pattern distributions can be correlated with the variables in question. In the example population in Fig. 4, variation along PC 1 correlates most strongly with average running speed on each trial, whereas variation along PC 2 correlates most strongly with experiment time. Running speed exerts different effects on the population code depending on experiment time; population code statistics also depend on the trial index even at a fixed running speed, reflecting slow changes in the internal state. This simultaneous variability implies that no single parameter fully determines the population state; rather, parameters influence the code in a non-additive, multiplexed manner. Across all analyzed populations, experiment time (i.e., trial index) and running speed correlate most strongly with the principal components that span the variability of spike pattern distributions. Trial-averaged pupil area and eye position show substantially lower correlation with code variation on individual trial timescales (Fig. 4l).

The population code may also be modulated by unobserved factors that are captured by changes in latent variable statistics. We therefore identified latent variables whose time averages correlate most strongly with positions along PC 1 and PC 2 (denoted latent variables A and B, respectively; see Fig. 4k). In the example case, latent variable A appears related to running speed, whereas latent variable B most likely reflects modulation by experiment time. In general, latent variable averages correlate more strongly with PC coordinates than behavioral parameters do (Fig. 4m), highlighting their potential to reveal additional factors modulating population codes during behavior.

### Ethological variability impacts the structure of the population code

Substantial variability in the dLGN population code implies that downstream circuits (e.g., in V1) receive nonstationary inputs even when repeatedly exposed to the same stimuli. To highlight these non-stationary changes in population activity, we first visualize how population synchrony changes along PC 1 (as shown in Fig. 4f; see Fig. 5a). We then select groups of 20 trials (corresponding to 10% of the total trial count) that are maximally separated along PC 1. Trials within each group can be further sorted according to their coordinates along PC 2 (Fig. 5b). While the similarity of population rasters within the same group is immediately apparent, this sorting reveals more nuanced variation. These effects are more clearly visible when comparing average spiking probabilities in each trial group sorted along PC 2 (Fig. 5c). In both groups, spiking probability follows a qualitatively similar pattern resembling the average change along PC 2 (Fig. 4h), even though overall firing rates and population synchrony are much higher in group B (blue). These patterns further highlight the multidimensional variation of population codes.

**Figure 5:**
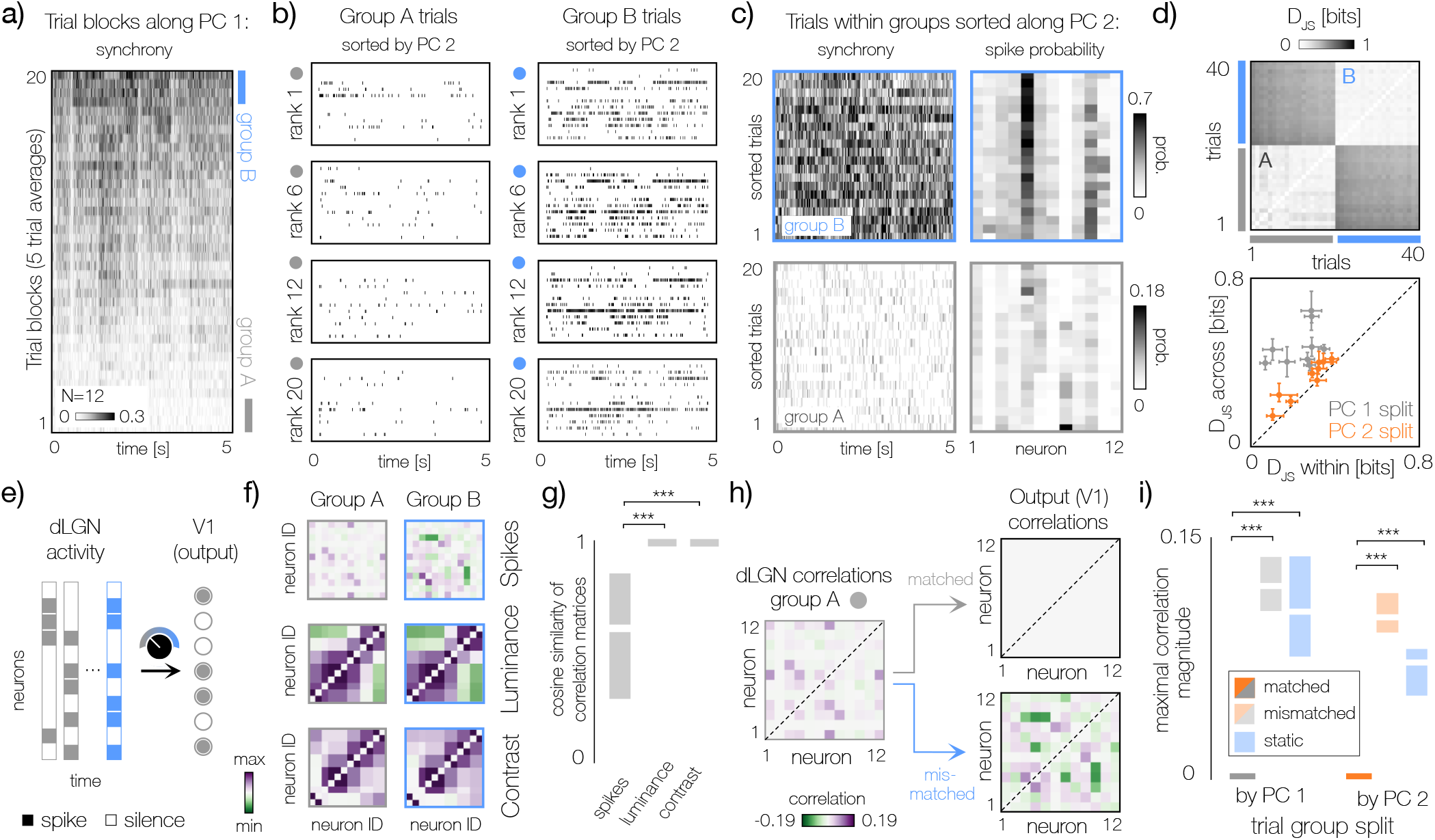
Ethological variability impacts the structure of the sensory code. **a)** Left: average synchrony in a population of 12 neurons (analyzed in Fig.4). Trials were sorted along the PC 1 and synchrony was computed by averaging blocks of 5 trials. **b)** Left: raster plots of individual trials from group A (with the lowest values of PC 1 coordinates), sorted along the PC 2 coordinate. Trial rank is denoted along the vertical axis. Right: same as A but for group B. **c)** Left: magnified trial groups marked with corresponding colors in a). Trial groups A and B consist of 20 trials with lowest and highest values of the coordinate along PC 1, respectively. Trials in both groups are sorted along PC 2. Right: average spiking probability on each trial within each group. **d)** Top: matrix of *D*_*JS*_ between trials in groups A and B. Bottom: dataset summary - *D*_*JS*_ across groups and within groups split along PC 1 (gray markers) and PC 2 (orange markers). Each symbol corresponds to a dataset. Dots denote to averages, bars denote standard deviations **e)** Downstream areas (e.g. V1) may need to adapt feed-forward connections to adapt to statistics of dLGN inputs that vary along different dimensions. **f)** Top: correlations between spikes in trial groups A and B split by PC 1. Middle: same as top but for local luminance within receptive field of each neuron. Bottom: same as middle but for local contrast. The color bar is rescaled differently for spike correlations (−0.2 to 0.2) and luminance/contrast correlations (−1 to 1). **g)** Dataset summary - cosine similarity between correlation matrices of spikes, local luminance and contrast computed for trial groups. Groups split along PC 1 and PC 2 were pooled together. Gray bars denote 25th-75th percentile, white lines denote median. Note that for stimulus luminance and contrast (middle and right bar respectively), the inter-quantile range is lower than the width of the rectangle. Stars denote statistical significance (paired sample t-test. For comparison spikes-luminance: *t* = −5.76, p-value = 2.30 × 10^−5^. For comparison spikes-contrast: *t* = −5.76, p-value = 2.31 × 10^−5^.) **h)** Correlations between neurons in the trial group A can be removed with a matched decorrelation transform (gray arrow). Application of a mismatched decorrelation transform optimized for group B (blue arrow) introduces spurious correlations. **i)** Dataset summary - maximal correlation magnitude after decorrelation. Splitting into trial groups along PC 1 and PC 2 is denoted by color (gray and orange, respectively). Matched and mismatched decorrelation transforms are denoted by saturated and faded colors, respectively. Decorrelation with a static transform fitted to all trials is denoted with blue. White lines denote median, and vertical color bars denote the 25th and 75th percentile across datasets and trial group splits. Stars denote statistical significance (for comparison with matched/mismatched: paired sample t-test, for PC 1 split - *t* = −13.50, p-value = 1.63 × 10^−10^, for PC 2 split - *t* = −13.36, p-value = 1.91 × 10^−10^. For comparison with static decorrelation: paired sample t-test, for PC 1 split - *t* = −11.99, p-value = 1.02 × 10^−10^, for PC 2 split - *t* = −12.48, p-value = 5.53 × 10^−10^).

We confirm the statistical separation of trial groups using the *D*_JS_ measure, which quantifies differences across entire spike pattern distributions without being limited to any specific feature of population activity (Fig. 5d, top). In all analyzed populations, trial groups that are maximally separated along PC 1 and PC 2 are statistically distinct and coherent, although, as expected, PC 2 generates weaker separation (Fig. 5d, bottom).

To understand the extent to which downstream circuits may need to adapt to changes in code statistics along each PC dimension, we must first specify a computational task. Here, we first consider decorrelation, a redundancy-reducing transformation known to maximize information transmission in the low-noise limit [46, 47]. In this setting, to effectively decorrelate spike patterns with non-stationary statistics, neurons in downstream areas (e.g., V1) must adapt their weights to the current population state (Fig. 5e) [29, 48].

We first observe that correlations between neurons are different in trial groups split along PC 1 (Fig. 5f, top row). To confirm that these differences do not result solely from changes in the sensory input caused by eye movements, we computed correlations between local luminance and contrast within each neuron’s receptive field, corrected for eye position (Fig. 5f, middle and bottom rows, respectively). Changes in neural correlations are much more substantial than in the inputs. This trend generalizes across all datasets, as well as to splits into trial groups along PC 1 and PC 2, as measured by the cosine similarity of correlation matrices (Fig. 5g). While correlations between stimulus features remain nearly unchanged across trial groups, correlations between spikes can differ dramatically. We note that here we consider correlations computed using all trials within a group, while in the decoding analyses (Fig. 4i) we computed correlations on each trial individually.

To simulate how downstream circuits could adapt to non-stationary correlations of dLGN populations, we implement a decorrelation transform using Zero-Phase Component Analysis (ZCA; see Methods). When the decorrelation transform is matched to each trial group, it effectively removes pairwise correlations between neurons and generates a non-redundant code (Fig. 5h, arrow marked “matched”). However, if the decorrelation transform is mismatched, it fails completely: not only does it not reduce reduce correlations between neurons, but it also introduces spurious dependencies (Fig. 5h, arrows marked “mismatched”). These effects generalize across all analyzed populations and extend to trial separations along PC 2 (Fig. 5i, gray and orange bars). To further test whether changes in the code structure necessitate adaptation by downstream circuits decorrelation, we performed a static decorrelation optimized for all trials (Fig. 5i, blue bars). The static decorrelation fails to remove redundancy from trial groups separated by PC 1 and PC 2, suggesting that an adaptive decorrelation transform is necessary. Overall, these findings imply that to maintain an efficient, decorrelated code, V1 circuits must dynamically adapt to the multidimensional variation of dLGN populations even when repeatedly exposed to the same sensory input.

### Ethological variability impacts the content of the population code

Transitions in population state may not only impact the statistical structure of the population code, but also modify the stimulus features that the code represents and conveys downstream, thus dynamically changing its content. One way to assess the content of the dLGN population code is to ask which stimulus frames can be distinguished from each other by downstream circuits, based on the patterns of dLGN activity they elicit [37, 41, 49]. As before, we rely on the *D*_JS_ dissimilarity between distributions of spike patterns. This time, however, we do not compare distributions of spike patterns generated on each trial. Instead, following previous work [34, 49], we compare distributions of spike patterns in response to different frames of the movie stimulus (Fig. 6a). If the dissimilarity equals 1 bit, these distributions are non-overlapping, and downstream neurons can reliably discriminate between the two stimulus frames that generated them. If it equals 0 bits, both stimulus frames generate identical distributions of spike patterns, making it impossible to tell them apart based on the population response.

**Figure 6:**
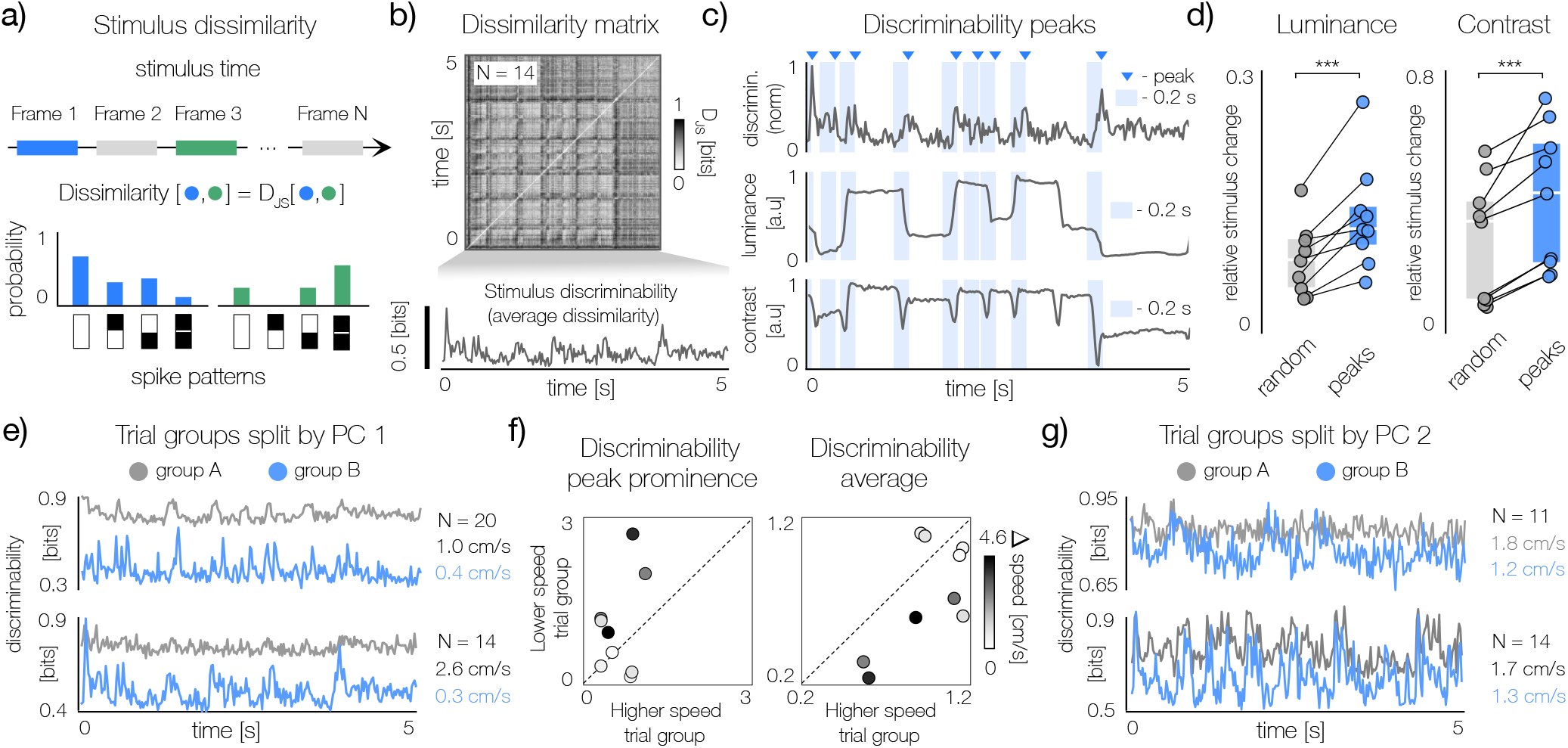
Ethological variability impacts the content of the sensory code. **a)** Stimulus dissimilarity is measured as *D*_*JS*_ between distributions of spike patterns elicited by two stimuli. **b)** Top: stimulus dissimilarity matrix for a population of 14 neurons. Bottom: stimulus discriminability computed by averaging dissimilarity per time-frame. **c)** Top: dissimilarity trace plotted in b) with peaks denoted by blue triangles. Blue shaded areas correspond to 200 ms interval immediately preceding each peak. Middle and bottom: stimulus luminance and contrast, respectively. Both signals were eye-position corrected and averaged based on outputs of receptive fields of individual neurons. **d)** Dataset summary - discriminability peaks around strong changes of the stimulus. Left: blue dots denote average relative change in luminance computed within 10 windows preceding each peak of discriminability. Gray dots denote randomly sampled stimulus intervals of the same length. Each pair of points corresponds to a dataset. White lines denote median and vertical gray bars denote 25th and 75th percentile of the distribution across datasets. Right: same as left but for the relative change in contrast. Stars denote statistical significance (paired sample t-test, for luminance - *t* = −4.93, p-value = 1.14 × 10^−3^, for contrast - *t* = −7.32, p-value = 8.25 × 10^−5^) **e)** Discriminability traces plotted for two example datasets: a population of 20 neurons (top) and a population of 14 neurons (bottom), where the same movie clip was used. Discriminability was computed for two trial groups split along PC 1. Blue and gray lines denote discriminability in the trial groups with lower and higher average running speed, respectively. **f)** Dataset summary - discriminability features in groups of trials split along PC 1. Each dot denotes a dataset. Grayscale denotes difference in the average running speed between two trial groups. Left: average prominence of discriminability peaks. Right: average discriminability. **g)** Top: stimulus discriminability computed for two trial groups split along PC 2 for a population of 11 neurons. Average running speed in each trial group is denoted by the color corresponding to the discriminability curve. Bottom: same as top but for a population of 14 neurons.

We computed the stimulus dissimilarity in different trial groups using the PRF model fit. The dissimilarity matrix expresses how distinguishable two stimulus frames are based on the population activity (Fig. 6b, top). Averaging the dissimilarity between a given stimulus frame and all other frames yields a measure of discriminability for that specific stimulus frame (Fig. 6b, bottom). A high discriminability value indicates that the frame can be easily distinguished from all other frames based on population responses. This quantity is reminiscent of stimulus-specific information (SSI), a measure of how informative an individual stimulus is by itself [50].

For all populations analyzed here, we identified trial groups that were maximally separated along PC 1 on the MDS plane (see Fig. 4). Next, for each of these groups, we computed stimulus discriminability for each movie frame (Fig. 6c, top row). When plotted as a function of stimulus time, discriminability tends to exhibit sharp peaks, indicating that certain stimulus frames are far more discernible than others based on dLGN population activity (Fig. 6c, top row, blue triangles). These peaks follow salient stimuli, i.e., strong relative changes in luminance and contrast (Fig. 6d). We note that stimulus luminance and contrast were computed based on eye-position-corrected receptive fields (see Methods).

To verify whether, in different states, dLGN populations prioritize accurate encoding of different stimulus features, we compared stimulus discriminability patterns in trial groups split along PC 1. We found particularly salient changes in discriminability between trial groups in datasets where PC 1 correlates strongly with the average running speed (Fig. 6e). In trial groups where the animal was mostly stationary, discriminability traces exhibited many sharp peaks corresponding to salient sensory events (Fig. 6e, blue lines), while other stimulus frames were only weakly discriminable. When the animal was running, discriminability substantially increased, suggesting an improvement in coding accuracy. This increase, however, was uniform across all stimulus frames and accompanied by the substantial reduction of peaks — a trend that generalizes across analyzed populations (Fig. 6f). Locomotion therefore triggers a transition not only in the overall precision of the sensory code (as reported previously for visual cortex [51, 52]), but also in the type of encoded stimuli. In a stationary animal, dLGN populations prioritize the encoding of salient sensory events, whereas during movement, stimuli are encoded with high accuracy regardless of saliency.

To complete the analysis, we also quantified stimulus discriminability in trial groups maximally separated along PC 2 (Fig. 6g). Although we observed salient examples of shifts in discriminability traces, we did not identify a general pattern. Overall, these results demonstrate that in the dLGN of a behaving animal, the content of the population code is also subject to substantial, multidimensional variability.

## DISCUSSION

Sensory ecology suggests that the nervous system evolved to efficiently process signals originating in the natural environment [53]. The underlying assumption is that neural activity across the sensory hierarchies is determined predominantly by external stimuli. However, the development of techniques to record neural activity in awake, behaving animals has revealed that sensory populations are constantly modulated by internal and behavioral states that influence sensory processing and inputs to higher stages of the visual brain [54]. This finding suggests that sensory pathways must adapt not only to statistics of the natural sensory world but also to the structure of the brain’s ethological variability during natural behavior.

In this work we approached this issue directly. Rather than focusing on individual modulatory factors such as running speed or arousal and attempting to study their impact in isolation, we characterized statistics of sensory messages conveyed by early sensory area dLGN during behavior. To this end we applied probabilistic raster factorization - a simple statistical model of population activity that was able to capture non-stationary dynamics of neuronal codes. We found that population responses in the dLGN of behaving mice are fundamentally non-stationary, with multiple processes contributing to this non-stationarity simultaneously. As a consequence, population codes can not only change their statistical structure, but also change their content, i.e., stimulus features they represent. This modulation results in multidimensional variability of population codes, which is crucially non-additive: the impact of multiple modulators cannot be reduced to a simple scalar increase in gain. Instead, it necessitates continuous adaptation to the changing structure and content of the sensory messages conveyed downstream by the visual thalamus.

### Challenges to ecological and stimulus-centric theories of sensory coding

Established theories of sensory processing assume that successive stages of sensory hierarchies process stimulus features of progressively increasing complexity to satisfy different objectives [55, 56]. These objectives may be general and task-agnostic, or focused on a specific task such as object recognition [2]. Regardless of focus, these theories share a common assumption: neural activity at each processing stage is largely determined by sensory signals from the external environment, rather than by internal signals arising within the brain. This assumption is directly challenged by contemporary studies of sensory coding during behavior.

A fundamental theoretical framework in sensory neuroscience, efficient coding, postulates that sensory neurons are adapted to the statistics of the signals they encounter in an organism’s ecological niche [1, 24]. Efficient coding theory has led to the development of the field of natural scene statistics, which seeks to explain properties of sensory systems by analyzing the structure of natural stimuli such as sounds or images [15]. Such analyses have led to major conceptual breakthroughs in our understanding of the objectives of sensory coding (e.g., [16, 57–59]) and perception (e.g. [60, 61]).

At its core, however, efficient coding theory is concerned with the statistics of inputs that neurons process, not necessarily with the statistics of natural scenes alone [62]. The profound non-stationarity and variability of sensory populations induced by internal state and behavior imply that the statistics of neural activity in sensory areas beyond the immediate periphery are only partially determined by the external environment. This challenges theories of sensory coding derived purely from external statistics of natural environments and calls for extending the study of natural scene statistics to the study of natural “sensory messages” conveyed to the brain by early sensory regions. For efficient coding to advance our understanding of sensory processing in behaving animals, it needs to account not only for sensory ecology but also for the ethology of natural behavior and fluctuating internal states.

Previous theoretical studies have presented elegant accounts of different forms of trial-to-trial variability as a form of adaptation to natural scene statistics, typically in the framework of perceptual inference (e.g., [22, 63, 64]). We foresee an exciting possibility of extending these approaches to account for the non-stationary structure of sensory codes in behaving animals.

A question of particular importance is whether downstream areas (e.g., V1) are “aware” of modulatory changes at the earlier stages of the visual hierarchy (e.g., dLGN) [65]. If modulation of sensory populations is driven by feedback connections, downstream areas that control this drive can immediately adapt to the state of the sensory code and thus can process inputs from lower regions accurately and efficiently. Recently proposed theories suggest that such feedback modulation may support tasks such as dynamic perceptual inference [26, 66], improve decoding accuracy [67], or adjust the sensory code to anticipated changes in stimulus statistics caused by animal movement [68]. The second possibility is that modulatory signals originate elsewhere and downstream areas receive inputs with constantly changing statistics, without awareness of when and how do these statistics change. Such a scenario is conceptually similar to sensory adaptation, where neurons need to infer a change in the distribution of a stimulus to maintain an efficient sensory code and to accurately perform necessary tasks [27–29]. These possibilities are non-exclusive and can co-exist. For example, feedback connections can influence a small population of neurons at a lower level of hierarchy, while other neurons may modulate their activity based on different factors.

Our analyses of the changes in the correlation structure of the dLGN suggest that downstream regions (e.g. V1) need to adapt their computations if their objective is to remove these correlations. Adaptive decorrelation has been observed empirically [69, 70] and studied theoretically [29]. The need for this computation, however, has been motivated by changing statistics of the sensory stimuli, not by fluctuations of internal states. Our results suggest that in order to maintain a decorrelated representation of dLGN activity, downstream neurons should also adapt to the structure of behavioral variability. A recently proposed framework for adaptive whitening of non-stationary signals suggests an intriguing solution to this problem [48].

### Computational function of the visual thalamus

Efficient coding theory has particularly strongly shaped our theoretical understanding of stimulus transformations implemented in the visual thalamus. In a now-classic study, Dan and colleagues showed that dLGN neurons in anesthetized cats temporally decorrelate natural video, thereby increasing coding efficiency [71]. Experimental work demonstrated further that the LGN selects only the most informative spikes and conveys them downstream to the visual cortex [72]. In this view, the LGN serves as a relay station that restructures retinal input into a more efficient representation by removing redundancies and transmitting this reformatted signal for further processing [73]. More recent work shows that LGN neurons in behaving rodents are subject to substantial behavioral modulation [11, 14] and are influenced by feedback connections [30]. The LGN may therefore be a particularly good starting point for extending the efficient coding framework to account not just for the statistics of natural scenes, but also for the multitude of internal states that modulate its activity.

Population coding in the thalamocortical system raises further general questions about hierarchical information processing in entirely natural conditions. In recent years, an increasing number of studies have highlighted the presence of movement-related and non-sensory signals in the visual cortex [74]. In mice, neurons across the visual hierarchy can be modulated by locomotion speed [3], head movements [75], and orofacial movements [9]. These behavioral signals may not only modulate the gain, but also impact the tuning of sensory neurons [76]. Beyond behavior, the visual cortex also encodes more abstract signals such as reward [77], task context [78, 79] or spatial position [80]. One of the experimental and conceptual challenges is to determine whether these multiplexed representations arise in the cortex or are already inherited from the earlier stages of the visual hierarchy [13]. Our results suggest the latter: at the population level, statistics of LGN inputs to the cortex already reflect concurrent modulatory processes.

### Extensions and future directions

To better understand how sensory systems interact with behavioral and internal states, we need to closely approximate not only ecological inputs but also ethological behaviors. In this study, we focused on analyses of neural recordings in a head-fixed animal, an experimental setup that is only a very coarse approximation of behavior in natural conditions. More recent experiments that enable recording of neural activity in freely behaving rodents [81] present an exciting opportunity to understand the structure and content of messages conveyed by neural populations in the wild.

A separate set of questions concerns the size of neural populations and the dimensionality of population codes. Here, we analyzed relatively small neural populations. Our approach, however, can be easily scaled to larger sets of neurons. This may be of particular importance when considering multidimensional variability of the population response statistics. The number of dimensions recovered by the manifold learning analysis, as well as the fraction of variance explained by PCA along each dimension, may depend entirely on the size of the recorded population and which neurons were recorded. Analyses of larger populations may therefore uncover additional states that modulate sensory codes during natural behavior.

## METHODS

### Population recordings

#### Mouse dLGN

The data was aquired in the Busse laboratory at LMU Munich and published before [30]. The experiments were designed to study the impact of optogenetic suppression of V1 feedback connections to LGN on neural coding. For analyses performed here we retained only trials where these feedback connections were not suppressed. We analyzed 9 population recordings from the dLGN of five mice (Fig. 1c, e right; see [30] for experimental details; see Supplementary Material for recording summary). The inclusion criterion was population size (larger than or equal to 10 neurons) and availability of the behavioral data. The animals were awake and head-fixed on a spherical treadmill. One of four 5-second natural movies was presented during each recording session; for each individual session, a single movie clip was used throughout. Movies were filmed close to the ground in wooded or grassy locations in Vancouver, BC to mimic animals’ views in their natural environment, and contained simulated gaze shifts, generated by manual camera movements.

#### Salamander retina

We used the data published in [32]. Spiking activity was recorded from the isolated retinal ganglion cell (RGC) layer of larval tiger salamander retinas during presentation of a 20-second natural movie stimulus (Fig. 1c, e; see [32] for experimental details).

#### Mouse retina

We used the data published in [33]. We used recordings from 8 isolated retina pieces from five mice, with a 25-second natural movie presented (Fig. 1c, e middle; see [33] for experimental details).

#### Data representation

We binned spike counts into bins of 20 ms (dLGN data) as well as 17 ms and 13 ms (salamander and mouse retinal data respectively). For all data used in this study, spike trains were binarized such that each time-frame was assigned a value of 1 if one or more spikes occurred, and 0 otherwise. The population response at each time-frame was represented as a binary vector 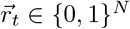, where *N* is the number of simultaneously recorded neurons and each entry *r*_*t,n*_ indicates whether the n-th neuron fired at least once within that bin (1 = spiking, 0 = silence).

### Eye position and pupil area extraction

The stimulus-viewing eye was illuminated with and infrared LED light (850 nm), and the eye was filmed with a camera (Guppy AVT camera; frame rate 50 Hz, Allied Vision, Exton, USA) equipped with a zoom lens (Navitar Zoom 6000). Eye videos were analyzed with DeepLabCut [82] using a 13-point eye model: 8 pupil markers evenly spaced around the pupil boundary, 2 eye corners (anterior, posterior), 2 eyelid points (upper, lower), and the corneal reflection. Markers with likelihood *<* 0.6 were discarded. Pupil boundaries were fit frame-by-frame as likelihood-weighted circles (*≥* 3 valid markers required), after rejecting spatial outliers among the remaining markers (points *>* 2× the median distance from the likelihood-weighted centroid). To get robust estimates of pupil position and size from per-frame fits, we applied a Kalman smoother on a 3-state constant-position model (pupil center x, y, radius). Unreliable segments, introduced by eye-closure, were removed according to pre-defined criteria. Pupil area was then computed from the smoothed radius (converted to *mm*^2^ using a corner-to-corner calibration distance of 3.25 mm), and the resulting time course was Gaussian-smoothed (*σ* = 250*ms*); Eye position was computed by rotating the pupil x/y into the coordinate frame defined by the two eye-corner markers, subtracting the corner-axis-defined eye center, and converting to visual angle (azimuth, elevation) via the arcsine of the displacement relative to eye radius derived from the measured corner-to-corner distance.

Some frames are dropped out because the pupil is covered by eye lid or the mouse is blinking. Those missing values are replaced with the trial median.

### Receptive field mapping and stimulus feature extraction

#### Receptive field fitting

Receptive fields (RFs) were mapped using sparse noise in a procedure described in [30]. For each position of the sparse noise squares (x, y), the envelope of multi-unit spiking activity (MUAe) of each electrode channel was averaged, normalized to a range between 0 and 1, and fitted with a two-dimensional Gaussian using the least-squares method. The receptive field was represented as a circle centered at (*x*_0_, *y*_0_) and described by the following parametric function 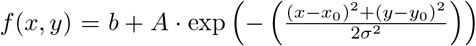 where b is the baseline activity, A is the peak activity, and *x*_0_ and *y*_0_ are the center coordinates of the receptive field. The parameter *σ* was fixed at 7.3^*°*^ for all units [83]. High-quality fits of receptive fields (*R*^2^ *>* 0.4) could be obtained for most channels in dLGN. For dLGN channels whose sampled visual field positions fell outside the sparse noise grid, or whose fits were of insufficient quality, receptive centers were estimated. For each session, separate linear regressions were fitted to the x and y coordinates of the well-fit receptive field centers as a function of channel depth, and these regressions were used to inter- or extrapolate the centers of the remaining channels. Each single unit was assigned the receptive field of the channel on which its extracellular waveform was largest. Of the 128 units in our dataset, 75 had well-fit measured receptive fields. The estimated receptive field center (*x*_0_, *y*_0_) was taken as the receptive field location at the mean eye position. At each time-frame t, the centers of receptive fields were adjusted to account for the instantaneous deviation of gaze in azimuth Δ*x*(*t*) and elevation Δ*y*(*t*) from the mean eye position.

#### Stimulus feature extraction

For each unit, the luminance and RMS contrast of the movies were extracted within its eye-position adjusted receptive field. Luminance was defined as the mean pixel intensity (0 to 255), and RMS contrast was defined as the standard deviation of the pixel intensity [84]. An adjusted luminance and contrast are computed with the adjusted RF center and a radius of 7.3 degrees surrounding it.

### Quantifying population response variability

To characterize trial-to-trial response variability, we computed the log-likelihood of each individual trial under the mean spiking probability. For each neuron *n* and time-frame *t*, the mean spiking probability was estimated by averaging binary responses across all *I* stimulus presentations: 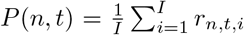. Assuming independent Bernoulli spiking, the log-likelihood of trial *i* takes the form: 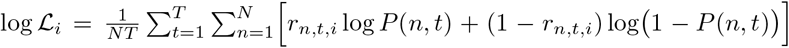 where *r*_*n,t,i*_ *∈ {*0, 1*}* is the binary spike response of neuron *n* at time-frame *t* on trial *i, N* is the number of neurons, and *T* is the number of time-frames.

### Probabilistic raster factorization model

The input to the PRF model is the binary matrix that represents population activity of *N* neurons over *T* time frames (note that these time frames can span multiple trials). The model learns *L* latent variables that approximate shared inputs to each neuron. The probability of a spike of neuron *n* in time bin *t* is defined as: 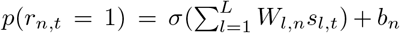, where *W*_*l,n*_ is the weight assigned by latent variable *l* to neuron *n, s*_*l,t*_ is the activation of that variable at time *t* and *b*_*n*_ is the neuron specific bias (see below). *σ* denotes the logistic function: 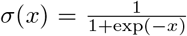. The bias term *b*_*n*_ is set such that the firing rate of each neuron is equal to it’s average across all data, when all latent variable activations are equal to 0. The bias term was set by hand for each neuron by computing the inverse-logit transform of that neurons average spiking probability. To regularize the model we apply a regularization term that minimizes deviations from the baseline spike probability for each neuron i.e. penalize the contributions of latent variables to the spiking probability in each bin. The resulting negative log-posterior (penalized log-likelihood) of the weight matrix *W* and activation matrix *S* takes the following form:

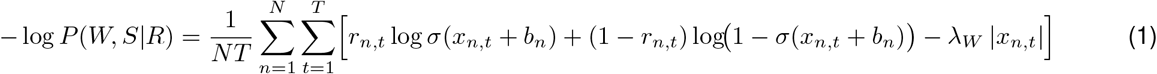

where 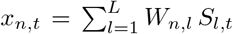 is the contribution of all latent variables to the spiking probability, *λ*_*w*_ is the regularization strength and *R* is the matrix of neural activity. The weight matrix *W* and latent coefficients *S* are fitted simultaneously by maximizing the log-likelihood via gradient descent, implemented using the MyGrad package in Python. The bias term *b*_*n,t*_ is fixed to the logit of the mean spiking probability for each neuron. The model was fitted separately for each dataset across *L ∈ {*1, …, *N}* latent variables (2c). To quantify the degree of explained uncertainty, in addition to the log-likelihood we computed the binary entropy for each fitted probability of individual spikes *p*(*r*_*n,t*_). The optimal *L* was selected as the smallest number of latent variables for which *ℒℒ* exceeded a predefined threshold (here −0.1 bits per neuron per time-frame).

#### Model evaluation

To assess how well the fitted model captures the statistics of the recorded population activity, we drew Monte Carlo samples of neural activity from the model fit. We then computed relevant statistics (spike probabilities, correlations, synchrony rasters and spike-pattern probabilities) and compared them against the data that the model was fitted to.

#### Raster marginal model

We computed the mean spiking probability for each neuron and time bin *p*(*r*_*n,t*_ = 1) averaged across trials. To generated a null distribution we sampled random trial rasters, where each spike had the probability determined by that average (Fig.2j).

#### Permutation test for latent–behavior correlations

To assess whether the correlation between each latent variable and each behavioral variable (running speed, pupil size, eye azimuth, eye elevation, and experiment time) exceeded chance, we constructed a null distribution by permutation. For each latent variable, the trial order was randomly permuted 5000 times, and the Pearson correlation with each behavioral variable was recomputed on each permutation. The significance threshold was set to the 95th percentile of the absolute null distribution, independently for each latent–behavior pair. A latent variable was considered significantly correlated with a behavioral variable if its observed |*ρ*| exceeded this threshold (Fig. 3d).

### Distance and dissimilarity measures

#### Jensen–Shannon divergence (*D*_JS_)

The Jensen-Shannon divergence *D*_JS_ between two distributions *P* and *Q* is defined as: 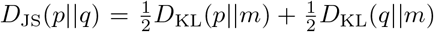 where 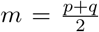 is the mixture distribution and 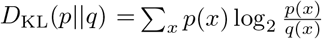 is the Kullback-Leibler divergence of distribution *q* from *p*. To evaluate the *D*_JS_ between population response distributions between trials, we approxinated distributions over spike patterns by generating Monte Carlo samples from the PRF model fit. We sampled each trial 1000 times from the model-estimated spiking probability resulting in 252000 samples per trial. We then computed the *D*_JS_ directly from the empirical distribution of population spike patterns estimated in that way.

#### Trial shuffles

To construct a null distribution for *D*_*JS*_, we performed Monte Carlo sampling by generating 1000 random datasets of the size of the original experiment. For each sample, the trial order of the model-predicted spiking probabilities was randomly permuted before drawing binary spike patterns, thus breaking the correspondence between the spiking probability profile and trial identity while preserving the marginal firing statistics of each neuron. The resulting null *D*_*JS*_ distribution reflects the expected pairwise divergence under the assumption that all trials share the same codeword distribution (Fig. 4d).

#### Euclidean distance between firing rates and correlations

For comparison with the Jensen-Shannon divergence, we computed the Euclidean distance between the spiking probabilities (firing rates) and correlation matrices computed for each trial individually. We denote them as *D*_FR_ and *D*_corr_ respectively.

### Manifold learning

#### Multidimensional scaling (MDS)

We applied multi-dimensional scaling to embed the pairwise *D*_JS_ distance matrix of each dataset into 2 to 10 dimensions. This analysis was implemented with the `manifold.MDS` function from the scikit-learn package in Python. We then computed the two principal components of the 2D MDS embedding of the *D*_JS_ distance matrix. PC1 is the axis capturing the most variance across trials; PC2 is the orthogonal axis capturing the remaining variance. This analysis was implemented with the `decomposition.PCA` function from the scikit-learn package in Python.

#### Decoding of a position on the MDS plane

For each PC of the MDS embedding (PC1 and PC2), we trained a linear decoder (ordinary least-squares regression) on 80% of trials and evaluated it on the held-out 20%. Two feature sets were compared as inputs to the decoder: (1) the mean spiking probability (firing rate) per neuron, averaged across time-frames within the trial (*N* features, where *N* is the number of neurons in the population); and (2) the upper triangle of the per-trial neuron–neuron Pearson correlation matrix (*N*(*N* − 1)*/*2 features), computed from binary spike data with a small jitter (distributed as a Gaussian with standard deviation of 10^−8^) added to avoid numerical issues. We performed cross-validation with *n* = 20 random 80/20 train/test splits, and quantified decoding performance as the Pearson correlation between the predicted and true PC scores on the test set.

## Decorrelation

We sorted the trials in descending order by PC1 score (Fig. 5a), and selected the first and last 10% trials to form two groups with maximally distinct spike pattern distributions. For each trial group, we concatenated all trials and computed the neuron–neuron correlation matrix. We performed Zero-phase Component Analysis (ZCA) whitening [85] to remove correlations and equalize variance of resulting outputs. This transformation can be thought of as a linear feed-forward neural network that yields a de-correlated representation of each inputs, and in which each output neuron is independent from its neighbors. Prior to ZCA activity of each neuron was z-scored (i.e. transformed to have the zero mean and unit standard deviation). To assess whether the correlation structure generalizes across trial groups, we also applied each group’s whitening matrix to the other group’s data (mismatched condition) and examined the residual correlations in the output (Fig. 5f). As a static transform we used a ZCA matrix fitted to all trials in the dataset concatenated together.

### Analysis of the code content

#### Stimulus dissimilarity

We measured stimulus dissimilarity between stimulus frames as the Jensen-Shannon divergence between spike pattern distributions in time bins corresponding to these stimulus frames within the trial. We denote these time bins as *t*_1_ and *t*_2_. The dissimilarity was then computed as 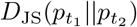. where 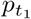 and (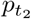 denotes population spike-pattern distributions in time bins *t*_1_ and *t*_2_ respectively. We approximated these distributions in trial groups via Monte Carlo sampling. For each time point we randomly drawn a trial corresponding to that group and generated random a spike pattern from the corresponding PRF fit. This procedure was used to generate the empirical distribution of spike patterns which we then used to approximate *D*_JS_ dissimilarity measure.

#### Discriminability

The stimulus discriminability *D* was measured by averaging the dissimilarity between the spike pattern distribution at *t* and those at all other time-frames: 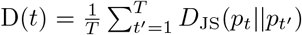 where *T* is the total number of time-frames in the trial. We identified the ‘peaks’ of the discriminability trace using the `signal.find peaks` function from the SciPy package in Python. Peaks were defined as time-frames with prominence *>* 0.2 and at least 5 time-frames away from neighboring peaks, where prominence measures how much a value stands out from the surrounding baseline of the signal, defined as the vertical distance between the peak value and its lowest contour line (Fig. 6b). After identifying peaks, we extracted 10 time-frames preceding each peak’s onset through the peak itself, for a total of 11 time-frames (220 ms; Fig. 6c). We computed the relative change of the stimulus associated with the *k*-th peak: 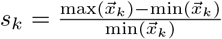, where 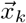 is the vector of stimulus contrast or luminance values within the sequence preceeding the *k* th peak. Contrast and luminance were estimated from receptive fields of each neuron individually (see above) and averaged after correcting for the eye position. For comparison, we established a null distribution by randomly sampling luminance and contrast sequences of the same length (11 time-frames, 220 ms) 1000 times for each trial group, and computing the relative change within these random sequences (Fig. 6d). Peak prominence plotted in Fig.6f was computed for each discriminability trace *D* as 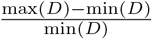.

## SUPPLEMENTARY MATERIAL

### Recording summary

We analyzed population activity from 9 recordings across 5 animals. Table 1 summarizes the number of simultaneously recorded neurons, number of trials, and fitted latent variables for each recording under a given log-likelihood threshold.

**Table 1:** Summary of recordings across datasets. Each row corresponds to one recording session, identified by animal id and recording id. *N*_neuron_ and *N*_trial_ denote the number of simultaneously recorded neurons and number of repetitions, respectively. *N*_latent_ denotes the number of latent variables selected for each recording, and *LL* denotes the corresponding log-likelihood threshold.

| Animal id | Recording id | $N_{\text{neuron}}$ | $N_{\text{trial}}$ | $N_{\text{latent}}, LL = -0.1$ | $N_{\text{latent}}, LL = -0.05$ |
| --- | --- | --- | --- | --- | --- |
| 2018-01 | 1 | 10 | 200 | 5 | 6 |
|  | 2 | 11 | 100 | 5 | 6 |
|  | 3 | 11 | 197 | 5 | 6 |
| 2019-02 | 1 | 14 | 200 | 5 | 6 |
| 2019-07 | 1 | 12 | 199 | 3 | 5 |
|  | 2 | 17 | 200 | 4 | 5 |
| 2019-08 | 1 | 14 | 200 | 5 | 6 |
|  | 2 | 14 | 200 | 3 | 4 |
| 2020-01 | 1 | 20 | 200 | 5 | 6 |

